# Benchmarks for Measurement of Duplicate Detection Methods in Nucleotide Databases

**DOI:** 10.1101/085324

**Authors:** Qingyu Chen, Justin Zobel, Karin Verspoor

**Affiliations:** Department of Computing and Information Systems The University of Melbourne, Parkville, VIC 3010, Australia

## Abstract

Duplication of information in databases is a major data quality challenge. The presence of duplicates, implying either redundancy or inconsistency, can have a range of impacts on the quality of analyses that use the data. To provide a sound basis for research on this issue in databases of nucleotide sequences, we have developed new, large-scale validated collections of duplicates, which can be used to test the effectiveness of duplicate detection methods. Previous collections were either designed primarily to test efficiency, or contained only a limited number of duplicates of limited kinds. To date, duplicate detection methods have been evaluated on separate, inconsistent benchmarks, leading to results that cannot be compared and, due to limitations of the benchmarks, of questionable generality.

In this study we present three nucleotide sequence database benchmarks, based on information drawn from a range of resources, including information derived from mapping to Swiss-Prot and TrEMBL. Each benchmark has distinct characteristics. We quantify these characteristics and argue for their complementary value in evaluation. The benchmarks collectively contain a vast number of validated biological duplicates; the largest has nearly half a billion duplicate pairs (although this is probably only a tiny fraction of the total that is present). They are also the first benchmarks targeting the primary nucleotide databases. The records include the 21 most heavily studied organisms in molecular biology research. Our quantitative analysis shows that duplicates in the different benchmarks, and in different organisms, have different characteristics. It is thus unreliable to evaluate duplicate detection methods against any single benchmark. For example, the benchmark derived from Swiss-Prot mappings identifies more diverse types of duplicates, showing the importance of expert curation, but is limited to coding sequences. Overall, these benchmarks form a resource that we believe will be of great value for development and evaluation of the duplicate detection methods that are required to help maintain these essential resources.

*Availability*: The benchmark data sets are available at https://bitbucket.org/biodbqual/benchmarks.

## 1. Introduction

Sequencing technologies are producing massive volumes of data. GenBank, one of the primary nucleotide databases, increased in size by over 40% in 2014 alone (1). However, researchers have been concerned about the underlying data quality in biological sequence databases since the 1990s (2). A particular problem of concern is duplicates, when a database contains multiple instances representing the same entity. Duplicates introduce redundancies, such as repetitive results in database search (3), and may even represent inconsistencies, such as contradictory functional annotations on multiple records that concern the same entity (4). Recent studies have noted duplicates as one of five central data quality problems (5), and it has been observed that detection and removal of duplicates is a key early step in bioinformatics database curation (6).

Existing work has addressed duplicate detection in biological sequence databases in different ways. This work falls into two broad categories: *efficiency-focused* methods that are based on assumptions such as that duplicates have identical or near-identical sequences, where the aim is to detect similar sequences in a scalable manner; and *quality-focused* methods that examine record fields other than the sequence, where the aim is accurate duplicate detection. However, the value of these existing approaches is unclear, due to the lack of broad-based, validated benchmarks; as some of this previous work illustrates, there is a tendency for investigators of new methods to use custom-built collections that emphasize the kind of characteristic their method is designed to detect.

Thus different methods have been evaluated using separate, inconsistent benchmarks (or test collections). The efficiency-focused methods used large benchmarks. However, the records in these benchmarks are not necessarily duplicates, due to use of mechanical assumptions about what a duplicate is. The quality-focused methods have used collections of expert-labelled duplicates. However, as a result of the manual effort involved, these collections are small and contain only limited kinds of duplicates from limited data sources. To date, no published benchmarks have included duplicates that are explicitly marked as such in the primary nucleotide databases, GenBank, the EMBL European Nucleotide Archive, and the DNA DataBank of Japan. (We refer to these collectively as *INSDC:* the International Nucleotide Sequence Database Collaboration (7).)

In this study, we address these issues by accomplishing the following:

- We introduce three benchmarks containing INSDC duplicates that were collected based on three different principles: records merged directly in INSDC (111,826 pairs); INSDC records labelled as references during Swiss-Prot expert curation (2,465,891 pairs); and INSDC records labelled as references in TrEMBL automatic curation (473,555,072 pairs);
- We quantitatively measure similarities between duplicates, showing that our benchmarks have duplicates with dramatically different characteristics, and are complementary to each other. Given these differences, we argue that it is insufficient to evaluate against only one benchmark; and
- We demonstrate the value of expert curation, in its identification of a much more diverse set of duplicate types.

It may seem that, with so many duplicates in our benchmarks, there is little need for new duplicate detection methods. However, the limitations of the mechanisms that led to discovery of these duplicates, and the fact that the prevalences are so very different between different species and resources, strongly suggest that these are a tiny fraction of the total that is likely to be present. While a half billion duplicates may seem like a vast number, they only involve 710,254 records, while the databases contain 189,264,014^1^ records altogether to date. Also, as suggested by the effort expended in expert curation, there is a great need for effective duplicate detection methods.

## 2. Background

In the context of general databases, the problems of quality control and duplicate detection have a long history of research. However, this work has only limited relevance for bioinformatics databases, because, for example, it has tended to focus on tasks such as ensuring that each real-world entity is only represented once, and the attributes of entities (such as ‘home address’) are externally verifiable. In this section we review prior work on duplicate detection in bioinformatics databases. We show that researchers have approached duplicate detection with different assumptions. We then review the more general duplicate detection literature, showing that the issue of a lack of rigorous benchmarks is a key problem for duplicate detection in general domains and to motivate our work. Finally, we describe the data quality control in INSDC, Swiss-Prot, and TrEMBL, as the sources for construction of the duplicate benchmark sets that we introduce.

### 2.1 Kinds of duplicate

Different communities, and even different individuals, may have inconsistent understandings of what a duplicate is. Such differences may in turn lead to different strategies for de-duplication.

A generic definition of a duplicate is that it occurs when there are multiple instances that point to the same entity. Yet this definition is inadequate; it requires a definition that allows identification of which things are ‘the same entity’. We have explored definitions of duplicates in other work (8). We regard two records as duplicates if, in the context of a particular task, the presence of one means that the other is not required. Here we explain that duplication has at least four characteristics, as follows.

First, duplication is not simply redundancy. The latter can be defined using a simple threshold. For example, if two instances have over 90% similarity, they can arguably be defined as redundant. Duplicate detection often regards such examples as ‘near duplicates’ (9) or ‘approximate duplicates’ (10). In bioinformatics, ‘redundancy’ is commonly used to describe records with sequence similarity over a certain threshold, such as 90% for CD-HIT (11). Nevertheless, instances with high similarity are not necessarily duplicates, and vice versa. For example, curators working with human pathway databases have found records labelled with the same reaction name that are not duplicates, while legitimate duplicates may exist under a variety of different names (12). Likewise, as we present later, nucleotide sequence records with high sequence similarity may not be duplicates, whereas records whose sequences are relatively different may be true duplicates.

Second, duplication is context dependent. From one perspective, two records might be considered duplicates while from another they distinct; one community may consider them duplicates whereas another may not. For instance, amongst gene annotation databases, more broader duplicate types are considered in Wilming et al. (13) than in Williams et al. (14), whereas, for genome characterization, ‘duplicate records’ means creation of a new record in the database using configurations of existing records (15). Different attributes have been emphasized in the different databases.

Third, duplication has various types with distinct characteristics. Multiple types of duplicates could be found even from the same perspective (8). By categorizing duplicates collected directly from INSDC, we have already found diverse types: similar or identical sequences; similar or identical fragments; duplicates with relatively different sequences; working drafts; sequencing in progress records; and predicted records. The prevalence of each type varies considerably between organisms. Studies on duplicate detection in general performance on a single dataset may be biased if we do not consider the independence and underlying stratifications (16). Thus, as well as creating benchmarks from different perspectives, we collect duplicates from multiple organisms from the same perspectives.

We do not regard these discrepancies as shortcomings or errors. Rather, we stress the diversity of duplication. The understanding of ‘duplicates’ may be different between database staff, computer scientists, biological curators, and so on, and benchmarks need to reflect this diversity. In this work we assemble duplicates from three different perspectives: expert curation (how data curators understand duplicates); automatic curation (how automatic software without expert review identifies duplicates); and merged-based quality checking (how records are merged in INSDC). These different perspectives reflect the diversity: a pair considered as duplicates from one perspective may not be so in another. For instance, nucleotide coding records might not be duplicates strictly at the DNA level, but they might be considered to be duplicates if they concern the same proteins. Use of different benchmarks derived from different assumptions tests the generality of duplicate detection methods: a method may have strong performance in one benchmark but very poor in another; only being verified from different benchmarks can possibly guarantee the method is robust.

Currently, understanding of duplicates via expert curation is the best approach. Here ‘expert curation’ means that curation either is purely manually performed, as in ONRLDB (17); or not entirely manual but involving expert review, as in UniProt/Swiss-Prot (18). Experts use experience and intuition to determine whether a pair is duplicate, and will often check additional resources to ensure the correctness of a decision (16). Studies on clinical (19) and biological databases (17) have demonstrated that expert curation can find a greater variety of duplicates, and ultimately improves the data quality. Therefore, in this work we derive one benchmark from UniProt/Swiss-Prot expert curation.

### 2.2 Impact of duplicates

There are many types of duplicate, and each type has different impacts on use of the databases. Approximate or near duplicates introduce redundancies, whereas other types may lead to inconsistencies.

Approximate or near duplicates in biological databases is not a new problem. We found related literature in 1994 (3), 2006 (20) and as recently as 2015.^2^ A recent significant issue was proteome redundancy in UniProtKB/TrEMBL (2015). UniProt staff observed that many records were over-represented, such as 5.97 million entries for just 1,692 strains of Mycobacterium tuberculosis. This redundancy impacts sequence similarity searches, proteomics identification, and motif searches. In total 46.9 million entries were removed.

Additionally, recall that duplicates are not just redundancies. Use of a simple similarity threshold will result in many false positives (distinct records with high similarity) and false negatives (duplicates with low similarity). Studies show that both cases matter: in clinical databases, merging of records from distinct patients by mistake may lead to withholding of a treatment if one patient is allergic but the other is not (21); failure to merge duplicate records for the same patient could lead to a fatal drug administration error (22). Likewise, in biological databases, merging of records with distinct functional annotations might result in incorrect function identification; failing to merge duplicate records with different functional annotations might lead to incorrect function prediction. One study retrieved corresponding records from two biological databases, Gene Expression Omnibus and ArrayExpress, but surprisingly found the number of records to be significantly different: the former has 72 whereas only 36 in latter. Some of the records were identical, but in some cases records were in one but not the other (23). Indeed, duplication commonly interacts with inconsistency (5).

Further, we cannot ignore the propagated impacts of duplicates. The above duplication issue in TrEMBL not only impacts TrEMBL itself, but also significantly impacts databases or studies using TrEMBL data. For instance, release of Pfam, a curated protein family database, was delayed for close to 2 years; the duplication issue in TrEMBL was the major reason (24). Even removal of duplicates in TrEMBL caused problems: ‘the removal of bacterial duplication in UniProtKB (and normal flux in protein) would have meant that nearly all (>90%) of Pfam seed alignments would have needed manual verification (and potential modification)… This imposes a significant manual biocuration burden’ (24).

Finally, duplicate detection across multiple sources provides valuable record linkages (25-27). Combination of information from multiple sources could link literature databases, containing papers mentioning the record; gene databases; and protein databases.

### 2.3 Duplicate detection methods

Most duplicate detection methods use pairwise comparison, where each record is compared against others in pairs using a similarity metric. The similarity score is typically computed by comparing the specific fields in the two records. The two classes of methods that we previously introduced, efficiency-focused and quality-focused, detect duplicates in different ways; we now summarize those approaches.

#### 2.3.1 Efficiency-focused methods

Efficiency-focused methods have two common features. One is that they typically rest on simple assumptions, such as that duplicates are records with identical or near-identical sequences. These are near or approximate duplicates as above. The other is application of heuristics to filter out pairs to compare, in order to reduce the running time. Thus, a common pattern of such methods is to assume that duplicates have sequence similarity greater than a certain threshold. In one of the earliest methods, *nrdb90,* it is assumed that duplicates have sequence similarities over 90%, with *k-mer* matching used to rapidly estimate similarity (28). In *CD-HIT,* 90% similarity is assumed, with short-substring matching as the heuristic (11); in *starcode*, a more recent method, it is assumed that duplicates have sequences with a Levenshtein distance of no more than 3, and pairs of sequences with greater estimated distance are ignored (29).

Using these assumptions and associated heuristics, the methods are designed to speed up the running time, which is typically the main focus of evaluation (11,28). While some such methods consider accuracy, efficiency is still the major concern (29). The collections are often whole databases, such as the NCBI non-redundant database^3^ for nucleotide databases and Protein Data Bank^4^ for protein databases. These collections are certainly large, but are not validated, that is, records are not known to be duplicates via quality-control or curation processes. The methods based on simple assumptions can reduce redundancies, but recall that duplication is not limited to redundancy: records with similar sequences may not be duplicates and vice versa. For instance, GenBank records `gi:15029538` and `gi: 8516100` have only 68% local identity,^5^ but submitters or database staff have merged them.^6^ Therefore, records measured solely based on a similarity threshold are not validated and do not provide a basis for measuring the accuracy of a duplicate detection method, that is, the false positive or false negative rate.

#### 2.3.2 Quality-focused methods

In contrast to efficiency-focused methods, quality-focused methods tend to have two main differences: use of a greater number of fields; and evaluation on validated datasets. An early method of this kind compared the similarity of both metadata (such as description, literature, and biological function annotations) and sequence, and then used association rule mining (30) to discover detection rules. More recent proposals focus on measuring metadata using approximate string matching: Markov random models (31), shortest-path edit distance (32), or longest approximately common prefix matching (33), the former two for general bioinformatics databases and the latter specifically for biomedical databases. The first method used a 1300-record dataset of protein records labelled by domain experts, whereas the others used a 1900-record dataset of protein records labelled in UniProt Proteomes, of protein sets from fully sequenced genomes in UniProt.

The collections used in this work are validated, but have significant limitations. First, both of the collections have less than 2,000 records, and only cover limited types of duplicates (47). We classified duplicates specifically on one of the benchmarks (merge-based) and it demonstrates that different organisms have dramatically distinct kinds of duplicate: in *Caenorhabditis elegans*, the majority duplicate type is identical sequences, whereas in *Danio rerio* the majority duplicate type is of similar fragments. From our case study of GC content and melting temperature, those different types introduce different impacts: duplicates under the exact sequence category only have 0.02% mean difference of GC content compared with normal pairs in *Homo sapiens*, whereas another type of duplicates that have relatively low sequence identity introduced a mean difference of 5.67%. A method could easily work well in a limited dataset of this kind but not be applicable for broader datasets with multiple types of duplicates. Second, they only cover a limited number of organisms; the first collection had two and the latter had five. Authors of prior studies, such as Rudniy et al. (33), acknowledged that differences of duplicates (different organisms have different kinds of duplicate; different duplicate types have different characteristics) are the main problem impacting the method performance.

In some respects, the use of small datasets to assess quality-based methods is understandable. It is difficult to find explicitly labelled duplicates. Typically, for nucleotide databases, sources of labelled duplicates are limited. In addition, these methods focus on the quality and so are unlikely to use strategies for pruning the search space, meaning that they are compute intensive. These methods also generally consider many more fields and many more pairs than the efficiency-focused methods. A dataset with 5000 records yields over 12 million pairs; even a small data set requires a large processing time under these conditions.

Hence there is no large-scale validated benchmark, and no verified collections of duplicate nucleotide records in INSDC. However, INSDC contains primary nucleotide data sources that are essential for protein databases. For instance, 95% of records in UniProt are from INSDC.^7^ A further underlying problem is that fundamental understanding of duplication is missing. The scale, characteristics, and impacts of duplicates in biological databases remain unclear.

### 2.4 Benchmarks in duplicate detection

Lack of large-scale validated benchmarks is a problem in duplicate detection in general domains. Researchers surveying duplicate detection methods have stated that the most challenging obstacle is lack of ‘standardized, large-scale benchmarking data sets’ (34). It is not easy to identify whether new methods surpass existing ones without reliable benchmarks. Moreover, some methods are based on machine learning, which require reliable training data. In general domains, many supervised or semi-supervised duplicate detection methods exist, such as decision trees (35) and active learning (36).

The severity of this issue is illustrated by the only supervised machine-learning method for bioinformatics of which we are aware, which was noted above (30). The method was developed on a collection of 1,300 records. In prior work, we reproduced the method and evaluated against a larger dataset with different types of duplicates. The results were extremely poor compared with the original outcomes, which we attribute to the insufficiency of the data used in the original work (37).

We aim to create large-scale validated benchmarks of duplicates. By assembling understanding of duplicates from different perspectives, it becomes possible to test different methods in the same platform, as well as test the robustness of methods in different contexts.

### 2.5 Quality control in bioinformatics databases

To construct a collection of explicitly labelled duplicates, an essential step is to understand the quality control process in bioinformatics databases, including how duplicates are found and merged. Here we describe how INSDC and UniProt perform quality control in general and indicate how these mechanisms can help in construction of large validated collections of duplicates.

#### 2.5.1 Quality control in INSDC

Merging of records addresses duplication in INSDC. The merge may occur due to various reasons, including cases where different submitters adding records for on the same biological entities, or changes of database policies. We have discussed various reasons for merging elsewhere (8). Different merge reasons reflect the fact that duplication may arise from diverse causes. Figure 1 shows an example. Record AC034192 is merged with Record AC087090 in Apr 2002. By contrast the different versions of Record AC034 (version 2 in Apr 2000 and version 3 in May 2000) are just normal updates on the same record. Therefore we only collect the former.

**Figure 1.**
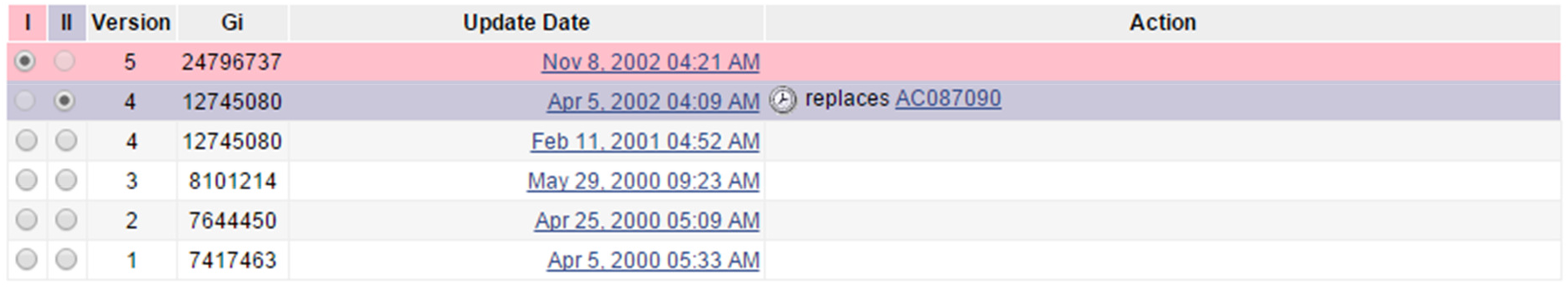
A screenshot of the revision history for record `gi:24796737`.^8^ Note the differences between normal updates (changes on a record itself) and merged records (duplicates). For instance, the record was updated from version 3 to 4, which is a normal update. A different record AC087090 is merged in during Apr 2002. This is a case of duplication confirmed by ENA staff. We only collected duplicates, not normal updates.

Staff confirmed that this is the only resource for merged records in INSDC. Currently there is no completely automatic way to collect such duplicates from the revision history. Elsewhere we have explained the procedure that we developed to collect these duplicates, why we believe that many duplicates are still present in INSDC, and why the collection is representative (8).

#### 2.5.2 Quality control in UniProt

UniProt Knowledgebase (UniProtKB) is a protein database that is a main focus of the UniProt Consortium. It has two sections: Swiss-Prot and TrEMBL. Swiss-Prot is expert curated and reviewed, with software support, whereas TrEMBL is curated automatically without review. Here we list the steps of curation in Swiss-Prot,^9^ as previously explained elsewhere (38):

1. Sequence curation: identify and merge records from same genes and same organisms; identify and document sequence discrepancies such as natural variations and frameshifts; explore homologs to check existing annotations and propagate other information;
2. Sequence analysis: predict sequence features using sequence analysis programs, then experts check the results;
3. Literature curation: identify relevant papers, read the full text and extract the related context, assign gene ontology terms accordingly;
4. Family curation: analyze putative homology relationships; perform steps 1-3 for identified instances;
5. Evidence attribution: link all expert curated data to the original source;
6. Quality assurance and integration: final check of finished entries and integration into Swiss-Prot.

Swiss-Prot curation is sophisticated and sensitive, and involves substantial expert effort, so the data quality can be assumed to be high. TrEMBL complements Swiss-Prot by using purely automatic curation. An example of automatic curation is use of expert-defined rules to determine protein functions and pathways.(39). Many instances of automatic curation correspond to the cross-references mapping to that record imported from nucleotide databases such as INSDC. INSDC records are cross-referenced during curation. The whole process is automatic and does not have expert review.

Therefore, it avoids expert curation with the trade-off of lower quality assurance. Overall both collections represent the state of the art in biological data curation.

Recall that nucleotide records in INSDC are primary sources for other databases. From a biological perspective, protein coding nucleotide sequences are translated into protein sequences (40). Both Swiss-Prot and TrEMBL cross-reference the coding sequence records in INSDC with their translated protein records. This provides a mapping between INSDC and curated protein databases. We can use the mapping between INSDC and Swiss-Prot and the mapping between INSDC and TrEMBL, respectively, to construct two collections of nucleotide duplicate records. We detail the methods and underlying ideas below.

## 3. Methods

We now explain how we construct our benchmarks, which we call the *merge-based, expert curation*, and *automatic curation* benchmarks; we then describe how we measure the duplicate pairs for all three benchmarks.

### 3.1 Benchmark construction

Our first benchmark is the *merge-based* collection, based on direct reports of merged records provided by record submitters, curators, and users to any of the INSDC databases. Creation of this benchmark involves searching the revision history of records in INSDC, tracking merged record IDs, and downloading accordingly. We have described the process in detail elsewhere, in work where we analyzed the scale, classification, and impacts of duplicates specifically in INSDC (8).

The other two benchmarks are the *expert curation* and *automatic curation* benchmarks. Construction of these benchmarks of duplicate nucleotide records is based on the mapping between INSDC and protein databases (Swiss-Prot and TrEMBL), and consists of two main steps. The first is to perform the mapping: downloading record IDs and using the existing mapping service; the second is to interpret the mapping results and find the cases where duplicates occur.

The first step has the following sub-steps. Our expert and automatic curation benchmarks are constructed using the same steps, except that one is based on mapping between INSDC and Swiss-Prot and the other is based on mapping between INSDC and TrEMBL.

1. Retrieve a list of coding records IDs for an organism in INSDC. We call these IIDs (I for INSDC). Databases under INSDC exchange data daily so the data is the same (though the representations may vary). Thus records can be retrieved from any one of the databases in INSDC. This list is used in the interpretation step;
2. Download a list of record IDs for an organism in either Swiss-Prot or TrEMBL. We call these UIDs (U for UniProt). This list is used in mapping;
3. Use the mapping service provided in UniProt (41) to generate mappings: Provide the UIDs from Step 2; Choose ‘UniProtKB AC/ID to EMBL/GenBank/DDBJ’ option; and Click ‘Generate Mapping’. This will generate a list of mappings. Each mapping contains the record ID in UniProt and the cross-referenced ID(s) in INSDC. We will use the mappings and IIDs in the interpretation step.

We interpret the mapping based on biological knowledge and database policies, as confirmed by UniProt staff. Recall that protein coding nucleotide sequences are translated into protein sequences. In principle, one coding sequence record in INSDC can be mapped into one protein record in UniProt; it can also be mapped into more than one protein record in UniProt. One gene may translate into multiple protein sequences. However, the reverse is not the case: one protein record can only be mapped into one nucleotide coding sequence record. The exception is that UniProt may merge multiple splice forms into a single protein record, each of them with a cross-reference to a nucleotide record. Here we assume the multiple splice forms represent a single record, since they are merged. More specifically, if one protein record in UniProt cross-references multiple coding sequence records in INSDC, those coding sequence records are duplicates. We classify the mappings into six cases, as follows.

- Case 1: A protein record maps to one nucleotide coding sequence record. No duplication is detected.
- Case 2: A protein record maps to many nucleotide coding sequence records. This is an instance of duplication. Here Swiss-Prot and TrEMBL represent different duplicate types. In the former splice forms, genetic variations, and other sequences are merged, whereas in the latter merges are mainly of records with close to identical sequences (either from the same or different submitters). That is also why we construct two different benchmarks accordingly.
- Case 3: Many protein records have the same mapped coding sequence records. There may be duplication, but we assume that the data is valid. For example, the cross-referenced coding sequence could be a complete genome that links to all corresponding coding sequences.
- Case 4: Protein records do not map to nucleotide coding sequence records. No duplication is detected.
- Case 5: The nucleotide coding sequences exist in IIDs but are not cross-referenced. Not all nucleotide records with a coding region will be integrated, and some might not be selected in the cross-reference process.
- Case 6: The nucleotide coding sequence records are cross-referenced, but are not in IIDs. A possible explanation is that the cross-referenced nucleotide sequence was predicted to be a coding sequence by curators or automatic software, but was not annotated as a coding sequence by the original submitters in INSDC. In other words, UniProt corrects the original missing annotations in INSDC. Such cases can be identified with the NOT_ANNOTATED_CDS qualifier on the DR line when searching in EMBL.

In this study we focus on Case 2, given that this is where duplicates are identified. We collected all the related nucleotide records and constructed the benchmarks accordingly.

### 3.2 Quantitative measures

After building the benchmarks as above, we quantitatively measured the similarities in nucleotide duplicate pairs in all three benchmarks to understand their characteristics. Typically, for each pair we measured the similarity of description, literature, and submitter, the local sequence identity, and the alignment proportion. The methods are described briefly here; more detail (Sections 3.2.1 to 3.2.3) is available in our other work(8).

#### 3.2.1 Description similarity

A description is provided in each nucleotide record’s DEFINITION field. This is typically a one-line description of the record, manually entered by record submitters. We have applied the following approximate string matching process to measure the description similarity of two records, using the Python *NLTK* package (42):

1. Tokenising: split the whole description word by word;
2. Lowering case: for each token, change all its characters into small cases;
3. Removing stop words: removes the words that are commonly used but not content bearing, such as ‘so’, ‘too’, ‘very’ and certain special characters;
4. Lemmatising: convert to a word to its base form. For example, ‘encoding’ will be converted to ‘encode’, or ‘cds’ (coding sequences) will be converted into ‘cd’;
5. Set representation: for each description, we represent it as a set of tokens after the above processing. We remove any repeated tokens;

We applied set comparison to measure the similarity using the Jaccard similarity defined by equation (1). Given two sets, it reports the number of shared elements as a fraction of the total number of elements. This similarity metric can successfully find descriptions containing the same tokens but in different orders.

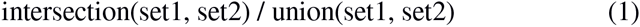

#### 3.2.2 Submitter similarity

The REFERENCE field of a record in the primary nucleotide databases contains two kinds of reference. The first is the literature citation that first introduced the record and the second is the submitter details. Here we measure the submitter details to find out whether two records are submitted by the same group.

We label a pair as ‘Same’ if it shares one of submission authors, and otherwise as ‘Different’. If a pair does not have such field, we label it as ‘N/A’. The author name is formatted as ‘last name, first initial’.

#### 3.2.3 Local sequence identity and alignment proportion

We used BLAST NCBI (version 2.2.30) (43) to measure local sequence identity. We used the *bl2seq* application that aligns sequences pairwise and reports the identity of every pair. NCBI BLAST staff advised on the recommended parameters for running BLAST pairwise alignment in general. We disabled the dusting parameter (which automatically filters low-complexity regions) and selected the smallest word size (4), aiming to achieve the highest accuracy as possible. Thus, we can reasonably conclude that a pair has low sequence identity if the output reports ‘no hits’ or the expected value is over the threshold.

We also used another metric, which we called the alignment proportion, to estimate the likelihood of the global identity between a pair. This has two advantages: in some cases where a pair has very high local identity, their lengths are significantly different. Use of alignment proportion can identify these cases; and running of global alignment is computationally intensive. Alignment proportion can directly estimate an upper bound on the possible global identity. It is computed using Formula (2) where L is the local alignment proportion, I is the locally aligned identical bases, D and R are sequences of the pair, and len(S) is the length of a sequence S.

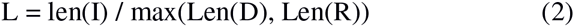

We constructed three benchmarks containing duplicates covering records for 21 organisms, using the above mapping process. We also quantitatively measured their characteristics in selected organisms. These 21 organisms are commonly used in molecular research projects and the NCBI Taxonomy provides direct links.^10^

**Table 1.** Submitter-based benchmark. Total records: numbers of records directly belong to the organism in total; Available merged groups: number of groups that are tracked in record revision histories. One group may contain multiple records. Duplicate pairs: total number of duplicate pairs. This table also appears in the paper (8)

| Organism | Total records | Available merged groups | Duplicate pairs |
| --- | --- | --- | --- |
| *Arabidopsis thaliana* | 337,640 | 47 | 50 |
| *Bos taurus* | 245,188 | 12,822 | 20,945 |
| *Caenorhabditis elegans* | 74,404 | 1,881 | 1,904 |
| *Chlamydomonas reinhardtii* | 24,891 | 10 | 17 |
| *Danio rerio* | 153,360 | 7,895 | 9,227 |
| *Dictyostelium discoideum* | 7943 | 25 | 26 |
| *Drosophila melanogaster* | 211,143 | 431 | 3,039 |
| *Escherichia coli* | 512,541 | 201 | 231 |
| *Hepatitis C virus* | 130,456 | 32 | 48 |
| *Homo sapiens* | 12,506,281 | 16,545 | 30,336 |
| *Mus musculus* | 1,730,943 | 13,222 | 23,733 |
| *Mycoplasma pneumoniae* | 1,009 | 2 | 3 |
| *Oryza sativa* | 108,395 | 6 | 6 |
| *Plasmodium falciparum* | 43,375 | 18 | 26 |
| *Pneumocystis carinii* | 528 | 1 | 1 |
| *Rattus norvegicus* | 318,577 | 12,411 | 19,295 |
| *Saccharomyces cerevisiae* | 68,236 | 165 | 191 |
| *Schizosaccharomyces pombe* | 4,086 | 39 | 545 |
| *Takifugu rubripes* | 51,654 | 64 | 72 |
| *Xenopus laevis* | 35,544 | 1,620 | 1,660 |

<sup>10</sup> http://www.ncbi.nlm.nih.gov/Taxonomy/taxonomyhome.html/

|  |  |  |  |
| --- | --- | --- | --- |
| *Zea mays* | 613,768 | 454 | 471 |

**Table 2.** Expert curation benchmark. Cross-referenced coding records: Number of records in INSDC that are cross-referenced in total; Cross-referenced coding records that are duplicates: Number of records that are duplicates based on interpretation of the mapping (Case 2); Duplicate pairs: total number of duplicate pairs.

| Organism | Cross-referenced coding records | Cross-referenced coding records that are duplicates | Duplicate pairs |
| --- | --- | --- | --- |
| *Arabidopsis thaliana* | 34,709 | 34,683 | 162,983 |
| *Bos taurus* | 9,605 | 5,646 | 28,443 |
| *Caenorhabditis elegans* | 3,225 | 2,597 | 4,493 |
| *Chlamydomonas reinhardtii* | 3,69 | 255 | 421 |
| *Danio rerio* | 5,244 | 3,858 | 4,942 |
| *Dictyostelium discoideum* | 1,242 | 1,188 | 1,757 |
| *Drosophila melanogaster* | 13,385 | 13,375 | 573,858 |
| *Escherichia coli* | 611 | 420 | 1,042 |
| *Homo sapiens* | 132,500 | 131,967 | 1,392,490 |
| *Mus musculus* | 74,132 | 72,840 | 252,213 |
| *Oryza sativa* | 4 | 0 | 0 |
| *Plasmodium falciparum* | 97 | 68 | 464 |
| *Pneumocystis carinii* | 33 | 19 | 11 |
| *Rattus norvegicus* | 15,595 | 11,686 | 24,000 |
| *Saccharomyces cerevisiae* | 84 | 67 | 297 |
| *Schizosaccharomyces pombe* | 3 | 3 | 2 |
| *Takifugu rubripes* | 153 | 64 | 59 |
| *Xenopus laevis* | 4,701 | 2,259 | 2,279 |
| *Zea mays* | 1,218 | 823 | 16,137 |

**Table 3.**
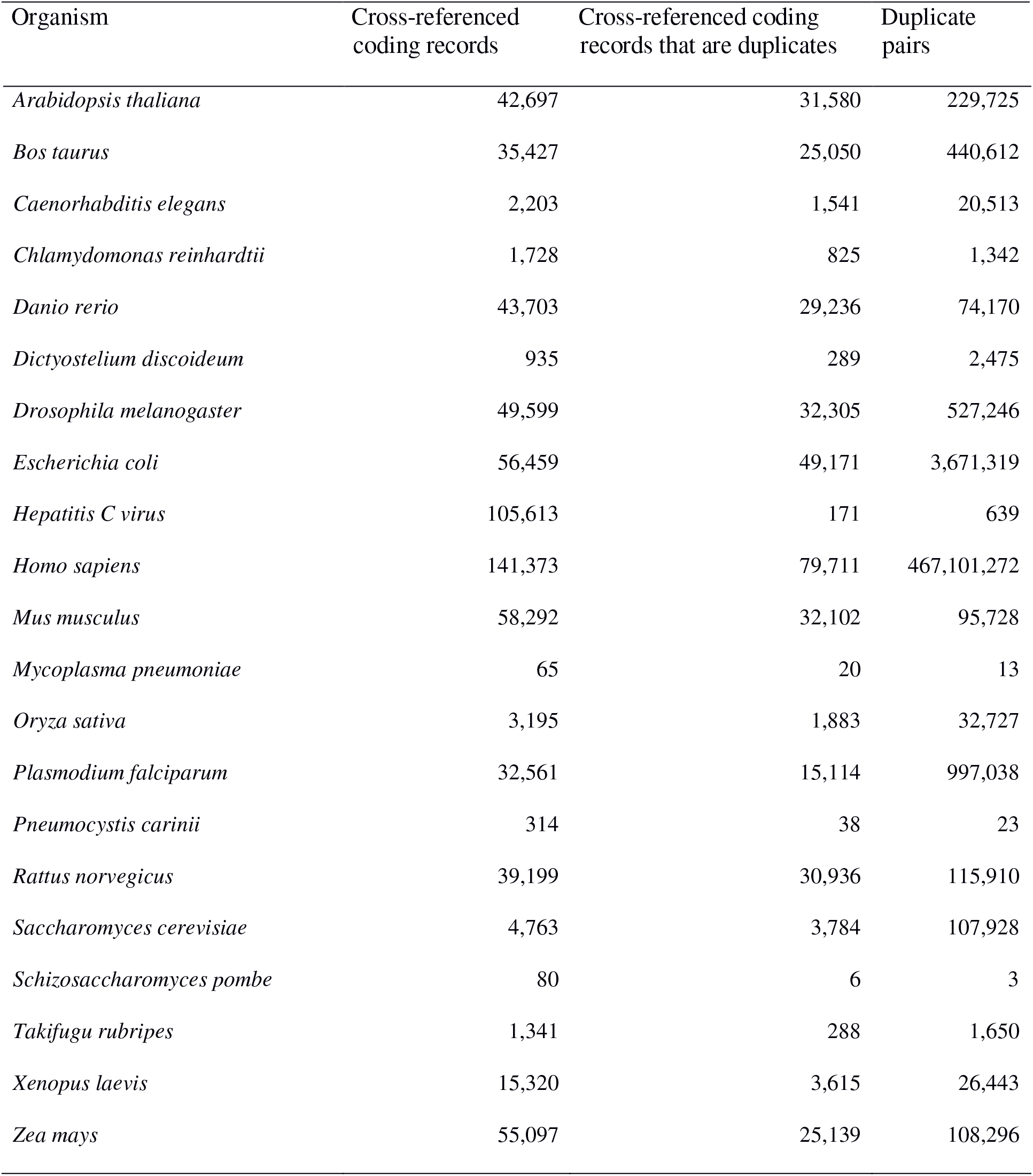
Automatic curation benchmark. The headings are the same as previously.

| Organism | Cross-referenced | Cross-referenced coding | Duplicate |
| --- | --- | --- | --- |

|  | coding records | records that are duplicates | pairs |
| --- | --- | --- | --- |
| *Arabidopsis thaliana* | 42,697 | 31,580 | 229,725 |
| *Bos taurus* | 35,427 | 25,050 | 440,612 |
| *Caenorhabditis elegans* | 2,203 | 1,541 | 20,513 |
| *Chlamydomonas reinhardtii* | 1,728 | 825 | 1,342 |
| *Danio rerio* | 43,703 | 29,236 | 74,170 |
| *Dictyostelium discoideum* | 935 | 289 | 2,475 |
| *Drosophila melanogaster* | 49,599 | 32,305 | 527,246 |
| *Escherichia coli* | 56,459 | 49,171 | 3,671,319 |
| *Hepatitis C virus* | 105,613 | 171 | 639 |
| *Homo sapiens* | 141,373 | 79,711 | 467,101,272 |
| *Mus musculus* | 58,292 | 32,102 | 95,728 |
| *Mycoplasma pneumoniae* | 65 | 20 | 13 |
| *Oryza sativa* | 3,195 | 1,883 | 32,727 |
| *Plasmodium falciparum* | 32,561 | 15,114 | 997,038 |
| *Pneumocystis carinii* | 314 | 38 | 23 |
| *Rattus norvegicus* | 39,199 | 30,936 | 115,910 |
| *Saccharomyces cerevisiae* | 4,763 | 3,784 | 107,928 |
| *Schizosaccharomyces pombe* | 80 | 6 | 3 |
| *Takifugu rubripes* | 1,341 | 288 | 1,650 |
| *Xenopus laevis* | 15,320 | 3,615 | 26,443 |
| *Zea mays* | 55,097 | 25,139 | 108,296 |

## 4. Results and Discussion

We present our results in two stages. The first introduces the statistics of the benchmarks constructed using the methods described above. The second provides the outcome of quantitative measurement of the duplicate pairs in different benchmarks.

We applied our methods to records for 21 organisms popularly studied organisms, listed in the NCBI Taxonomy website.^11^ Tables 2, 3, and 4 show the summary statistics of the duplicates collected in the three benchmarks. Table 2 is reproduced from another of our papers (8). All the benchmarks are significantly larger than previous collections of verified duplicates. The submitter-based benchmark has over 100,000 duplicate pairs. Even more duplicate pairs are in the other two benchmarks: the expert curation benchmark has around 2.46 million pairs and the automatic curation benchmark has around 0.47 billion pairs; hence these two are also appropriate for evaluation of efficiency-focused methods.

We measured duplicates for *Bos taurus, Rattus norvegicus, Saccharomyces cerevisiae, Xenopus laevis* and *Zea mays* quantitatively as stated above. Figures 1-8 show representative results, for *Xenopus laevis* and *Zea mays*. These figures demonstrate that duplicates in different benchmarks have dramatically different characteristics, and that duplicates from different organisms in the same benchmarks also have variable characteristics. We elaborate further as follows.

**Figure 2.**
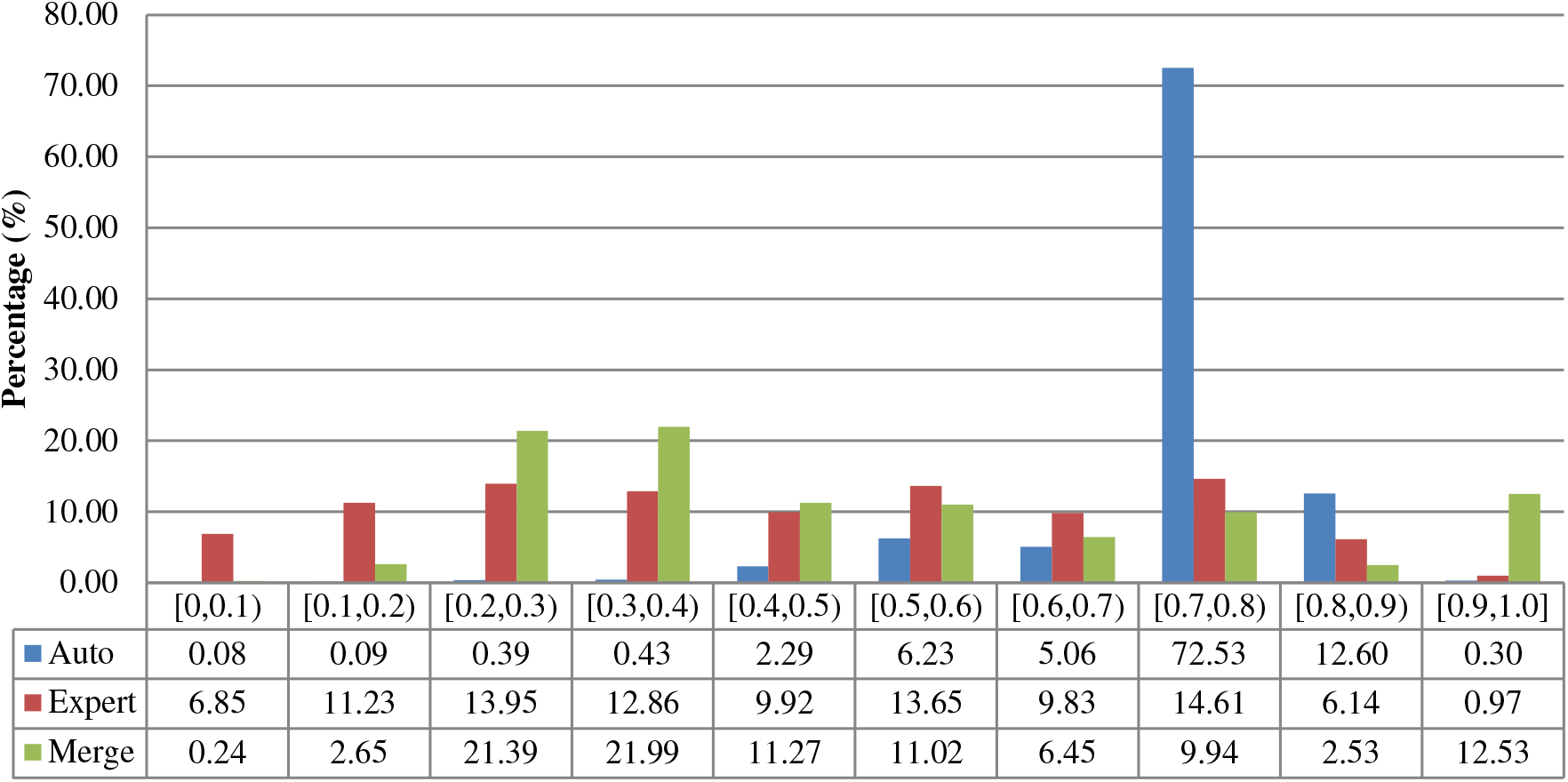
Description similarities of duplicates from *Xenopus laevis* in three benchmarks: Auto for auto curation based; Expert for expert curation; and Merge for merge-based collection. X-axis defines the similarity range. For instance, [0.5, 0.6) means greater than or equal to 0.5 and less than 0.6. Y-axis defines the proportion for each similarity range.

**Figure 3.**
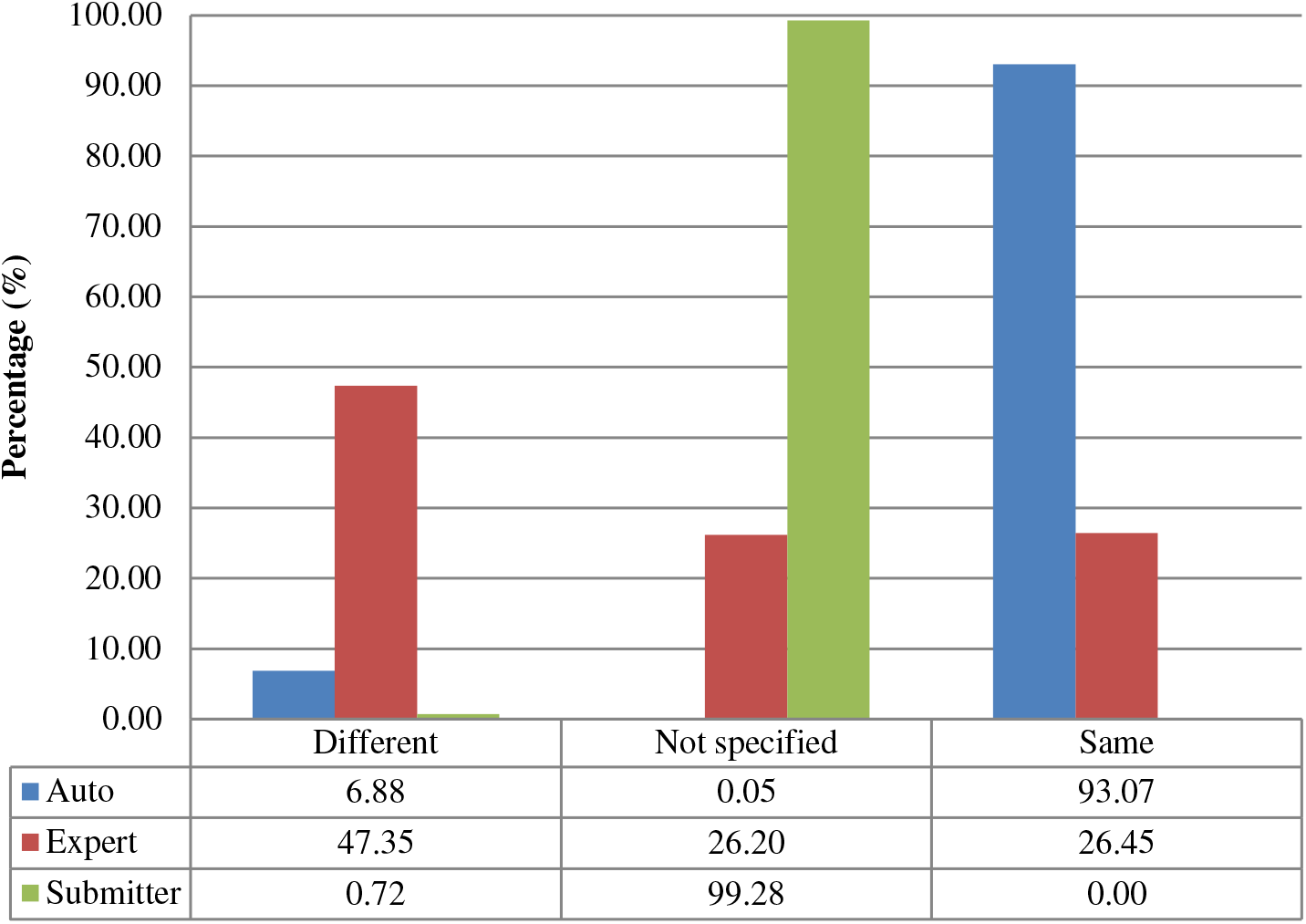
Submitter similarities of duplicates from *Xenopus laevis* in three benchmarks. Different: the submitters of records are completely Different; Same: the pair at least shares with at least one submitter; Not specified: no submitter details are specified in REFERENCE field in records by standard. The rest is the same as above.

**Figure 4.**
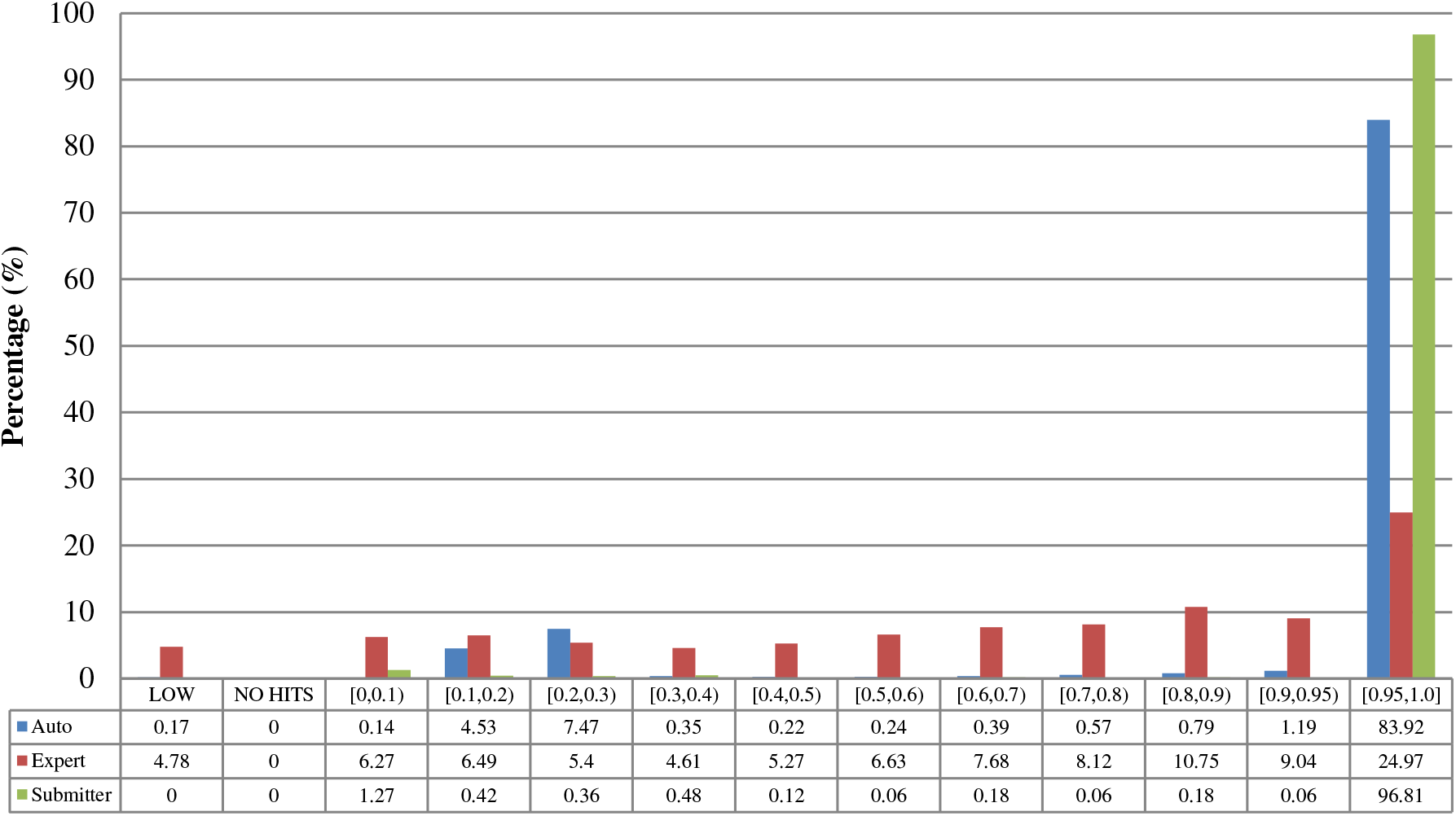
Alignment proportion of duplicates from *Xenopus laevis*. LOW refers to similarity that is greater than the threshold or NO HITS based on BLAST output. Recall that we chose the parameters to produce reliable BLAST output.

**Figure 5.**
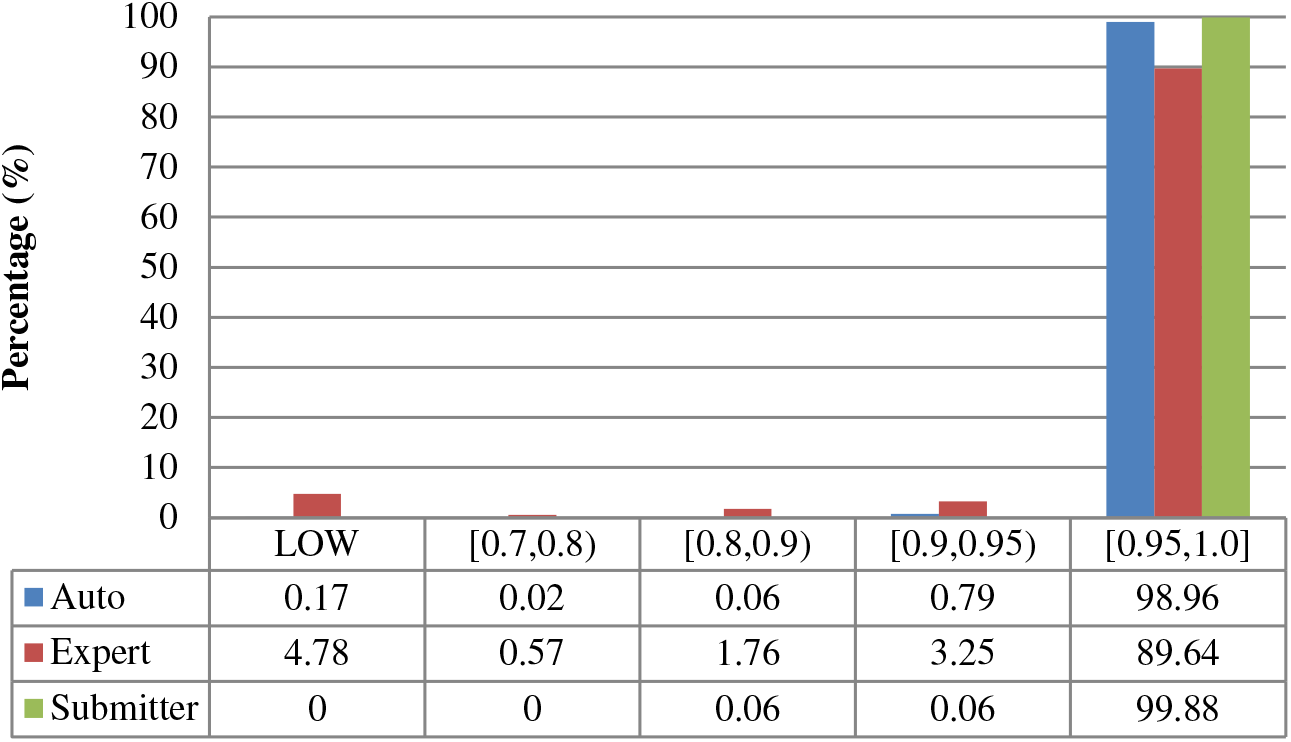
Local sequence identity of duplicates from *Xenopus laevis* in three benchmarks. The rest is the same as above.

**Figure 6.**
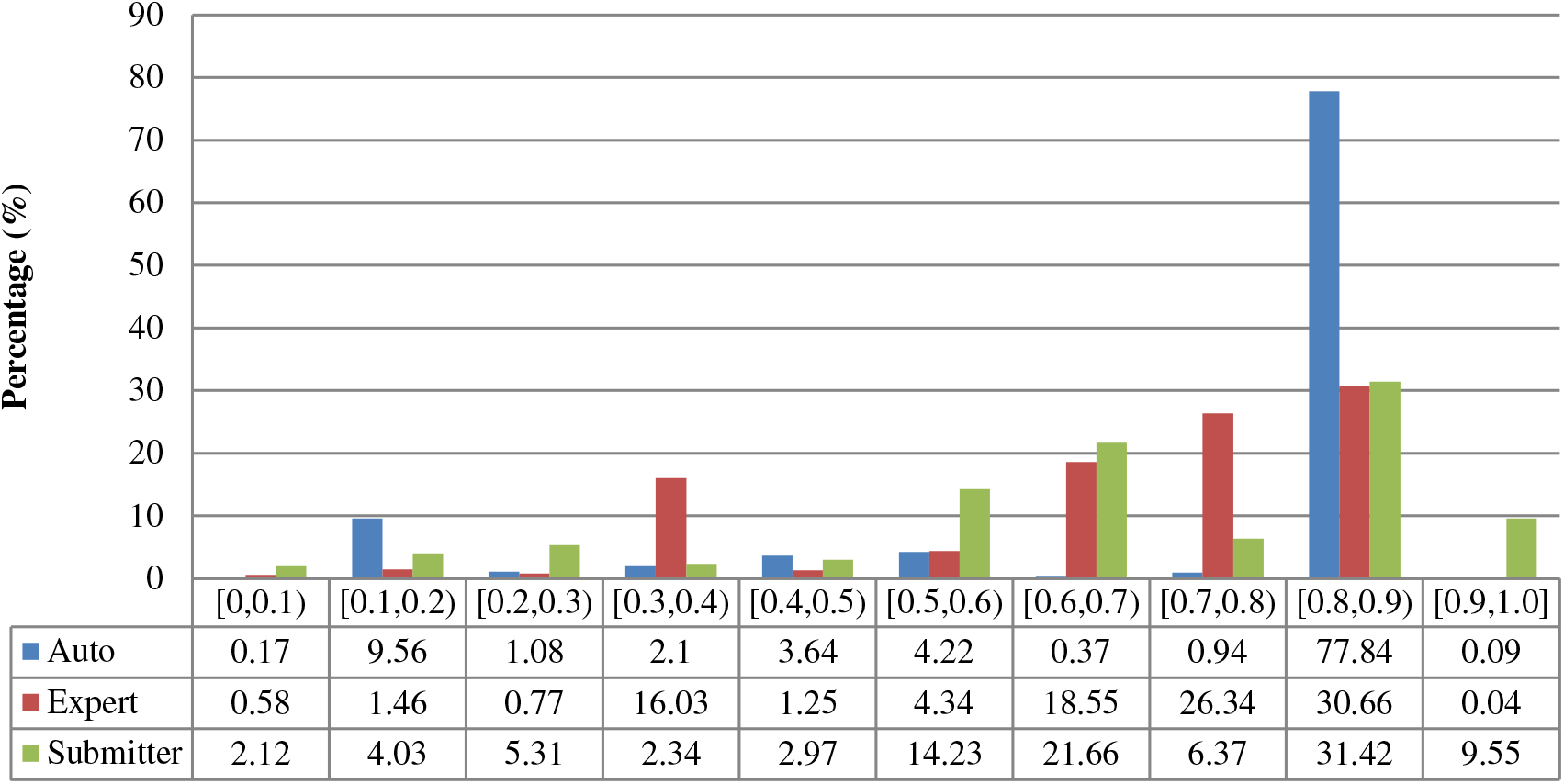
Description similarity of duplicates from *Zea mays* in three benchmarks.

**Figure 7.**
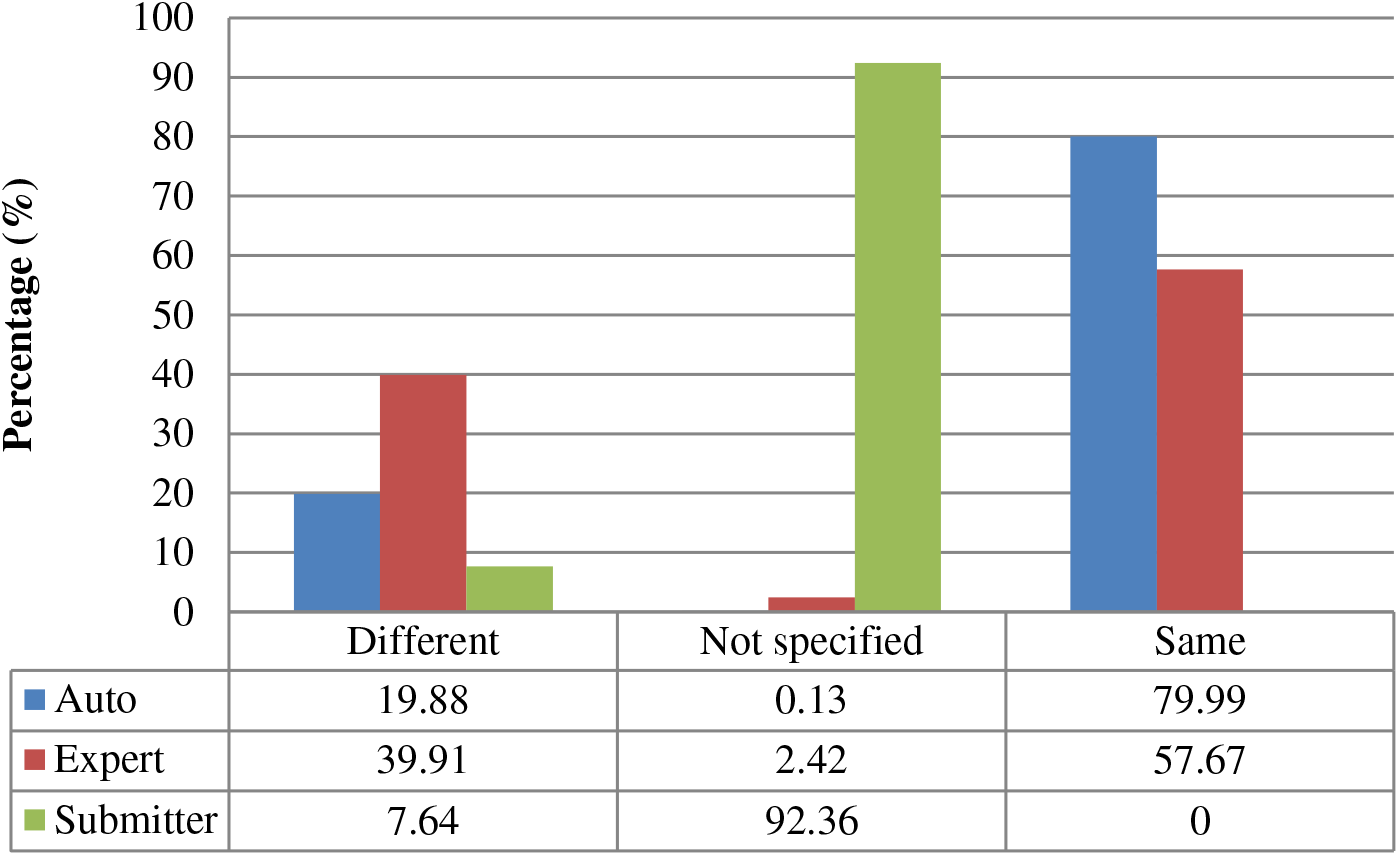
Submitter similarity of duplicates from *Zea mays* in three benchmarks.

**Figure 8.**
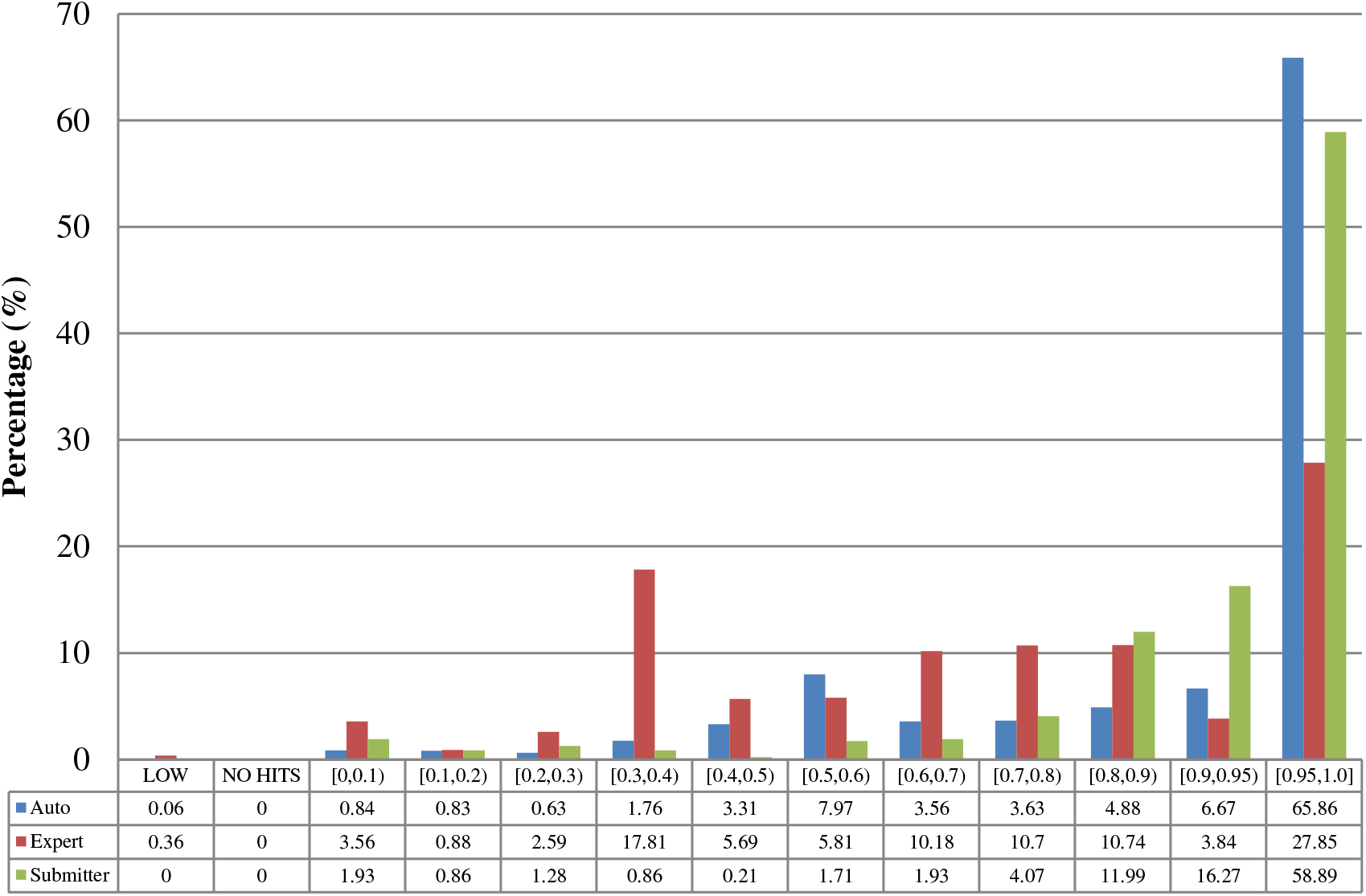
Alignment proportion of duplicates from *Zea Mays* in three benchmarks.

**Figure 9.**
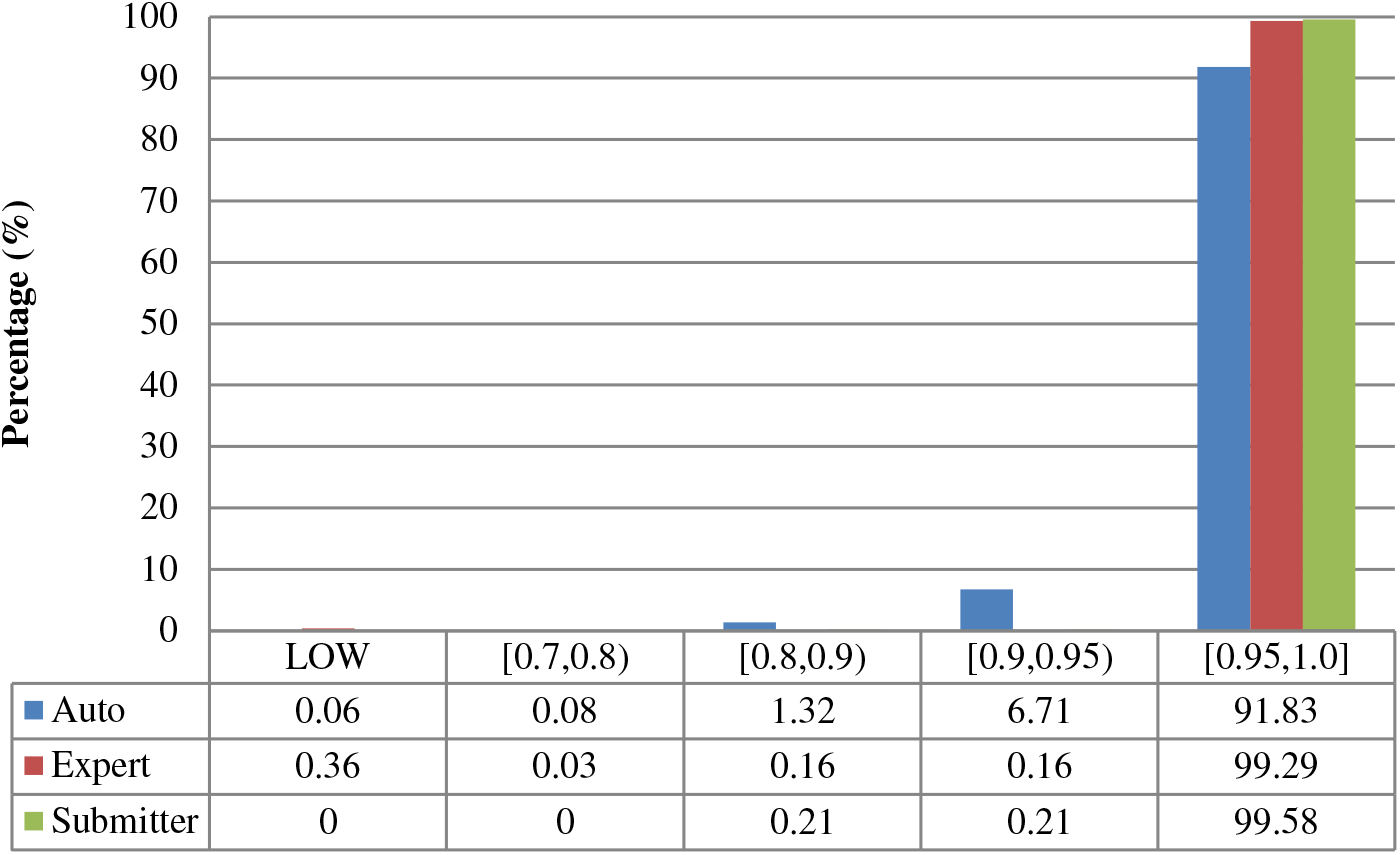
Local sequence identity of duplicates from *Zea Mays* in three benchmarks.

Construction of benchmarks from three different perspectives has yielded different numbers of duplicates with distinct characteristics in each benchmark. These benchmarks have their own advantages and limitations. We analyse and present them here.

- The merge-based benchmark is broad. Essentially all types of records in INSDC are represented, including clones, introns, and binding regions; all types in addition to the coding sequences that are cross-referenced in protein databases. However, only limited duplicates have been identified by this method. Our results clearly show that it contains far fewer duplicates than the other two, even though the original total number of records is much larger.
- The expert curation benchmark is shown to contain a much more diverse set of duplicate types. For instance, Figure 4 clearly illustrates that expert curation benchmark identifies much more diverse kinds of duplicate in *Xenopus Laevis* than the other two benchmarks. It not only identifies 25.0% of duplicates with close to the same sequences, but it finds presence of duplicates with very different lengths and even duplicates with relatively low sequence identity. In contrast, the other two mainly identify duplicates having almost the same sequence - 83.9% for automatic curation benchmark and 96.8% for the merge-based benchmark. However, the volume of duplicates is smaller than for automatic curation. The use of the protein database means that only coding sequences will be found.
- The automatic curation benchmark holds the highest number of duplicates amongst the three. However, even though it represents the state-of-the-art in automatic curation, it mainly uses rule-based curation and does not have expert review, so is still not as diverse or exhaustive as expert curation. For example, in Figure 1, over 70% of the identified duplicates have high description similarity, whereas the expert curation benchmark contains duplicates with description similarity in different distributions. As with the expert curation benchmark, it only contains coding sequences by construction.

The analysis shows that these three benchmarks complement each other. Merging records in INSDC provides preliminary quality checking across all kinds of records in INSDC. Curation (automatic and expert) provides more reliable and detailed checking specifically for coding sequences. Expert curation contains more kinds of duplicates and automatic curation has a larger volume of identified duplicates.

Recall that previous studies used a limited number of records with a limited number of organisms and kinds of duplication. Given the richness evidenced in our benchmarks, and the distinctions between them, it is unreliable to evaluate against only one benchmark, or multiple benchmarks constructed from the same perspective. As shown above, the expert curation benchmark contains considerable numbers of duplicates that have the distinct alignment proportions or relatively low similarity sequences. The efficiency-focused duplicate detection methods discussed earlier thus would fail to find many of the duplicates in our expert curation benchmark.

Also, duplicates in one benchmark yet in different organisms have distinct characteristics. For instance, as shown in figures for *Xenopus laevis* and *Zea mays,* duplicates in *Zea mays* generally have higher description similarity (comparing Figure 2 with Figure 6), submitted by more same submitters (comparing Figure 3 with Figure 7), more similar sequence lengths (comparing Figure 4 with Figure 8), and higher sequence identity (comparing Figure 4 with Figure 8). However, duplicates in *Xenopus laevis* have different characteristics. For instance, the expert curation benchmark contains 40.0% and 57.7% of duplicates submitted by different and same submitters respectively. Yet the same benchmark shows many more duplicates in *Xenopus laevis* from different submitters (47.4%), which is double the amount for the same submitters (26.4%). Due to these differences, methods that demonstrate good performance on one organism may not display comparable performance on others.

Additionally, the two curation-based benchmarks indicate that there are potentially many undiscovered duplicates in the primary nucleotide databases. Using *Arabidopsis thaliana* as an example, submitters or database staff (recall that both can request a record merge when noticing duplicates) only identified 47 groups of duplicates out of 337,640 records in total. The impression from this would be that the overall prevalence of duplicates in INSDC is quite low. However, Swiss-Prot and TrEMBL only cross-referenced 34,709 and 42,697 *Arabidopsis thaliana* records respectively, yet tracing their mappings results in finding that 34,683 (99.93%) records in Table 2 and 31,580 (73.96%) records in Table 3 have at least one corresponding duplicate record, even though they only examine coding sequences. It may be possible to construct another benchmark through the mapping between INSDC and RefSeq, using the approach described in this paper.

Another observation is that Swiss-Prot, with expert curation, contains a more diverse set of duplicates than the other benchmarks. From the results, it can be observed that expert curation can find occurrences of duplicates that have low description similarity, are submitted by completely different groups, have varied lengths, or are of comparatively low local sequence identity. This illustrates that it is not sufficient to focus on duplicates that have highly similar sequences of highly similar lengths. A case study has already found that expert curation rectifies errors in original studies (39). Our study on duplicates illustrates this from another angle.

These results also highlight the complexity of duplicates that are present in bioinformatics databases. The overlap amongst our benchmarks is remarkably minimal. The submitter benchmark includes records that do not correspond to coding sequences, so they are not considered by the protein databases. Swiss-Prot and TrEMBL use different curation processes as mentioned above. It shows that from the perspective of one resource, a pair may be considered as a duplicate, but on the basis of another resource may not be.

More fundamentally, records that are considered as duplicates for one task may not be duplicates for another. Thus it is not possible to use a simple and universal definition to conceptualize duplicates. Given that the results show that kinds and prevalence of duplicates vary amongst organisms and benchmarks, it suggests that studies are needed to answer fundamental questions: what kinds of duplicates are there? What are their corresponding impacts for biological studies that draw from the sequence databases? Can existing duplicate detection methods successfully find the type of duplicates that has impacts for specific kinds of biomedical investigations? These questions are currently unanswered. The benchmarks here enable such discovery (47). We explored the prevalence, categories, and impacts of duplicates in the submitter-based benchmark to understand the duplication directly in INSDC.

To summarise, we review the benefits of having created these benchmarks.

First, the records in the benchmarks can be uses for two main purposes: (1) as duplicates to merge; (2) as records to label or cross-reference to support record linkage. We now examine the two cases:

*Case 1*: record `gi:15029538`^12^ and `gi:8516100`^13^. This is an example that we noted earlier from the submitter collection. Record `gi:8616100` was submitted by the Whitehead Institute/MIT Center for Genome Research. It concerns the RP11-301H18 clone in *Homo sapiens* chromosome 9. It has 18 unordered pieces as the submitters documented. The later record `gi:15029538` was submitted by the Sanger Centre. That record also concerns the RP11-301H18 clone but it only has 3 unordered pieces. So this case shows an example of duplication where different submitters submit records about the same entities. Note that they are inconsistent, in that both the meta data and sequence are quite different. Therefore a merge was done (by either database staff or submitter). Record `gi:8616100` was replaced by `gi:15029538`, as `gi:15029538` has fewer unordered pieces, that is, is closer to being complete. Then record `gi:8616100` became obsolete. Only record `gi:15029538` can be updated. This record now has complete sequence (no unordered pieces) around 2012, after 18 updates from the version since the merge.

*Case 2:* record `gi:21263169`^14^, `gi:18490316`^15^, and `gi:7020685`^16^. These records are from the expert curation collection. At the protein level, they correspond to the same protein record Q8TBF5, about a Phosphatidylinositol-glycan biosynthesis class X protein. Those three records have been explicitly cross-referenced into the same protein entry during expert curation. The translations of record `gi:18490316` and `gi:7020685` are almost the same. Further, the expert-reviewed protein record Q8TBF5 is documented as follows^17^:

- AC055725 [`gi:21263169`] Genomic DNA. No translation available;
- BC022542 [`gi:18490316`] mRNA. Translation: AAH22542.1. Sequence problems;
- AK000529 [`gi:7020685`] mRNA. Translation: BAA91233.1. Sequence problems;

Those annotations were made via curation to mark problematic sequences submitted to INSDC. The ‘no translation available’ annotation indicates that the original submitted INSDC records did not specify the coding sequence (CDS) regions, but the UniProt curators have identified the CDS. ‘Sequence problems’ refers to ‘discrepancies due to an erroneous gene model prediction, erroneous ORF assignment, miscellaneous discrepancy, etc.’^18^ resolved by the curator. Therefore, without expert curation, it is indeed difficult to access the correct information and is difficult to know they refer to the same protein. As mentioned earlier, an important impact of duplicate detection is record linkage. Cross-referencing across multiple databases is certainly useful, regardless of whether the linked records are regarded as duplicates.

Second, considering the three benchmarks as a whole, they cover diverse duplicate types. The detailed types are summarized elsewhere (47), but broadly three types are evident: (1) similar records, if not identical; (2) fragments; (3) somewhat different records belonging to the same entities. Existing studies have already shown all of them have specific impacts on biomedical tasks. Type (1) may affect database searches (44); type (2) may affect meta-analyses (45); while type (3) may confuse novice database users.

Third, those benchmarks are constructed based on different principles. The large volume of the dataset, and diversity in type of duplicate, can provide a basis for evaluation of both efficiency and accuracy. Benchmarks are always a problem for duplicate detection methods: a method can detect duplicates in one dataset successfully, but may get poor performance on another. This is because the methods have different definitions of duplicate, or those datasets have different types or distributions. This is why the duplicate detection survey identified the creation of benchmarks as a pressing task (46). Multiple benchmarks enable testing of the robustness and generalization of the proposed methods. We used six organisms from the expert curated benchmark as the dataset and developed a supervised learning duplicate detection method (47). We tested the generality of the trained model as an example: whether a model trained from duplicate records in one organism maintains the performance in another organism. This is effectively showing how users can use the benchmarks as test cases, perhaps organized by organisms or by type.

## 5. Conclusion

In this study we established three large-scale validated benchmarks of duplicates in bioinformatics databases, specifically focusing on identifying duplicates from primary nucleotide databases (INSDC). The benchmarks are available for use at https://bitbucket.ors/biodbaual/benchmarks. These benchmark data sets can be used to support development and evaluation of duplicate detection methods. The three benchmarks contain the largest number of duplicates validated by submitters, database staff, expert curation, or automatic curation presented to date, with nearly half a billion record pairs in the largest of our collections.

We explained how we constructed the benchmarks and their underlying principles. We also measured the characteristics of duplicates collected in these benchmarks quantitatively, and found substantial variation among them. This demonstrates that it is unreliable to evaluate methods with only one benchmark. We find that expert curation in Swiss-Prot can identify much more diverse kinds of duplicates and emphasize that we appreciate the effort of expert curation due to its finer-grained assessment of duplication.

In future work, we plan to explore the possibility of mapping other curated databases to INSDC to construct more duplicate collections. We will assess these duplicates in more depth to establish a detailed taxonomy of duplicates and collaborate with biologists to measure the possible impacts of different types of duplicates in practical biomedical applications. However, this work already provides new insights into the characteristics of duplicates in INSDC, and has created a resource that can be used for the development of duplicate detection methods. With, in all likelihood, vast numbers of undiscovered duplicates, such methods will be essential to maintenance of these critical databases.

## Acknowledgements

We greatly appreciate the assistance of Elisabeth Gasteiger from Swiss-Prot, who advised on and confirmed the mapping process in this work with domain expertise. We also thank Nicole Silvester and Clara Amid from the EMBL European Nucleotide Archive, who advised on the procedures regarding merged records in INSDC. Finally we are grateful to Wayne Mattern from NCBI, who advised how to use BLAST properly by setting reliable parameter values. This work was supported by the ARC Discovery program, grant number DP150101550.

## Footnotes

1 http://www.ddbj.nig.ac.jp/breakdown_stats/dbgrowth-e.html#ddbjvalue

2 http://www.uniprot.org/help/proteome_redundancy

3 Listed at https://blast.ncbi.nlm.nih.gov/Blast.cgi?PAGE_TYPE=BlastSearch

4 http://www.rcsb.org/pdb/home/home.do

5 Measured in Section 3.2 advised by NCBI BLAST staff

6 http://www.ncbi.nlm.nih.gov/nuccore/AL592206.2?report=girevhist

7 http://www.uniprot.org/help/sequence_origin

8 http://www.ncbi.nlm.gov/nuccore/AC034192.5?report=girevhist

9 http://www.uniprot.org/help/

10 http://www.ncbi.nlm.nih.gov/Taxonomy/taxonomyhome.html/

11 http://www.ncbi.nlm.nih.gov/Taxonomy/taxonomyhome.html/

12 https://www.ncbi.nlm.nih.gov/nuccore/15029538

13 https://www.ncbi.nlm.nih.gov/nuccore/AC069109.2?report=genbank

14 https://www.ncbi.nlm.nih.gov/nuccore/AC055725

15 https://www.ncbi.nlm.nih.gov/nuccore/BC022542

16 https://www.ncbi.nlm.nih.gov/nuccore/AK000529

17 http://www.uniprot.org/uniprot/Q8TBF5

18 http://www.uniprot.org/help/cross_references_section

